# Accurate quantification of lysyl oxidase concentration in human tissue

**DOI:** 10.1101/2023.07.06.548012

**Authors:** Yimin Yao, Lara Perryman, Amna Zahoor, Ross Hamilton, Jessica Stolp, Wolfgang Jarolimek

## Abstract

The family of Lysyl oxidase enzymes play fundamental roles in the formation of the extracellular matrix, through catalyzing the crosslinking of collagen and elastin fibers. Lysyl oxidase (LOX) — one of the 5 family members (LOX, LOXL1-4), is a disease biomarker, with blood concentration positively correlating with progression of fibrosis or cancer. An accurate quantification of LOX concentration can support diagnosis, monitoring of disease progression or treatment success. However, reported LOX concentrations in human blood are inconsistent. Therefore, a novel, high-throughput and sensitive digital enzyme-linked immunosorbent assay was developed using two validated and selective human anti-LOX antibodies and single molecule array technology. Both, the 56 kDa pro-form and the 32 kDa active form can be accurately measured from recombinant and native protein. The serum LOX concentration correlated with LOX activity measured in the same platform using a bio-probe. The usefulness of this technology was demonstrated in serum from bladder cancer patients wherein LOX concentration was significantly higher compared to the healthy subjects. This study demonstrates the validation and use of a sensitive and accurate method for measuring LOX concentration in human samples. This novel method may be superior than some commercially available enzyme-linked immunosorbent assay kits for accurate measurement of LOX concentrations in clinical settings.

## Introduction

Lysyl oxidase (LOX) is a copper-dependent amine oxidase that belongs to a five-membered family of enzymes [LOX and LOX-like (LOXL) 1-4]. Like all LOX family members, LOX facilitates the formation and repair of the extracellular matrix by catalyzing oxidative deamination of lysine and hydroxylysine residues to aminoadipic semialdehydes which subsequently form covalent crosslinks of collagen and elastin (1). LOX is synthesized as a 56-kDa pro-form and released into the extracellular space where it is proteolytically cleaved by bone morphogenetic protein 1 and other procollagen C-proteinases to release the mature 32-kDa active form (2).

Serum and plasma LOX concentrations have been reported in a number of clinical studies showing their upregulation in a range of fibrotic diseases and cancers (S1 Table). As a result, LOX blood concentration could be a valuable biomarker to support the diagnosis and monitor progression of such diseases. It is accepted that the usefulness of a diagnostic biomarker rests on a reliable, accurate and high-throughput assay with a lower limit of quantification below normally occurring values. Furthermore, concordant results should be obtained from qualified sites and operators running the same biomarker test (3). However, reported serum LOX concentrations from healthy human subjects vary largely between studies and range from 0.2 ng/mL to > 40 ng/mL (4–10). This inconsistency in reported LOX concentrations may be due to the differences in commercial enzyme-linked immunosorbent assay (ELISA) kits supplied from different sources (S1 Table).

In addition to the ability to measure LOX concentration in liquid biopsies, LOX is also the predominantly expressed isoform of the lysyl oxidase family in the skin, providing an opportunity for an easily accessible tissue biopsy (11). The protein expression of LOX has been shown to be upregulated in fibrotic skin conditions such as scleroderma (12). To date, there is a lack of an assay that can accurately quantify the LOX protein level and activity in a small biopsy of human skin.

Based on a previous study in which a novel digital ELISA (DEA) has been developed to measure the concentration and activity of LOXL2 (13). The current study aims to develop and validate a high-throughput and sensitive protein concentration assay that accurately quantifies LOX concentration in human samples using high resolution single molecule array (SIMOA) technology (S1 Fig), and to adopt such a platform to measure LOX activity.

## Methods

### Ethics for human blood and tissue samples

Human serum was obtained from Bioivt & Elevating Science subjected to Institutional and Independent Review Board approval from the U.S. Food and Drug Administration. Among 19 patients with bladder cancer, sixteen patients were in stage 1, two patients were in stage 2 and another patient’s cancer stage unidentified (Ethics Application No: 05035, 15002 and 800962). The participants involved in the current study were recruited and samples taken during 07 January 2016 to 30 October 2019 for Ethics Application No: 05035, 06 February 2019 to 07 October 2019 for Ethics Application No: 15002 and on 03 March 2016 for Ethics Application No: 800962. Control serum samples were collected from healthy volunteers with no demographic information provided. Measurements were done by blinded analysis and no patient ID were recorded.

The use of human skin biopsies for the measurement of LOX concentration and target engagement was approved by Belberry as part of the Phase 1 clinical trials ACTRN12621000322831 (Ethics Application No: 2020-12-1296-A-3). The participants involved in the current study were recruited during March 2021 to May 2021, and skin biopsies were collected within 3 hours before receiving treatment at first post-screen visit during 30 March 2021 to 07 May 2021. Skin biopsies were stored immediately after collection at −80 °C until subsequent analysis for LOX activity and concentration. Measurements were done by blinded analysis and no patient ID were recorded.

### Production and extraction of lysyl oxidases

#### Native LOX production and extraction from fibroblast cell lines

NHLF (Lonza, CC-2512), IMR90 (ATCC, CCL-186) and BRL3A (Cellbank Australia, 85111503) were grown in their respective growth medium as per manufacturer’s instructions in humidified atmosphere of 5% CO_2_ and 95% air at 37°C. At 90% confluency, cells were washed twice with warm sterile phosphate buffered saline (PBS) (Lonza, BE17-516F) and incubated in starving medium [DMEM phenol red free (Thermo Fisher Scientific, 31053028), 0.1% fetal bovine serum (Thermo Fisher Scientific, A3161002), NEAA (Lonza, 13-114E) and 10 µM CuSO_4_]. Recombinant human (rh) TGF-β (10 ng/mL, R&D Systems, 7754-BH) was applied to NHLF cell line only to induce LOX expression. After 3 days, the conditioned medium (CM) was collected and filtered (0.45 µM). Concentration (∼40×) and buffer exchange to 6 M urea, 50 mM sodium borate buffer, pH 8.2 was performed with 10 kDa Amicon filters (Merck Millipore, ACS501024). The concentrated CM was further fractionated with 50-kDa Amicon filters (Merck Millipore, ACS505024), and the lower molecular weight portion, containing the 32 kDa LOX protein was collected.

#### Bovine aorta LOX extraction

Bovine aortas were collected at an abattoir minutes after the termination of 3-6 months old calves. The tunica intima and media were separated from the other layers, washed with PBS and blotted dry before being quickly frozen in dry ice and then stored at −80°C. For LOX extraction, 30 g of tissue was pulverized in liquid nitrogen and resuspended in cold wash buffer containing 0.15 M NaCl, 50 mM sodium borate, pH 8.0 with 0.25 mM PMSF (Sigma-Aldrich, P7626) and 1 µL/mL aprotinin (Sigma-Aldrich, A6279) supplemented as protease inhibitors. The mixture was homogenized (T25 IKA Turrax) for 2 minutes on ice then washed thrice with same volumes of ice-cold wash buffer. Between each wash, samples were briefly homogenized and centrifuged at 4500 *g* for 20 minutes at 4°C and supernatant discarded. The pellet was resuspended and kept in extraction buffer containing 6 M urea, 50 mM sodium borate, pH 8.2 with the same as protease inhibitors as above, overnight at 4°C. The mixture was then centrifuged at 10,000 *g* for 20 minutes at 4°C. The supernatant was further purified by incubating with 0.1 g/mL hydroxyapatite (Sigma-Aldrich, 21223) and Cleanascite (10% w/w, Biotech Support Group, NC0542680) for 30 minutes at 4°C, then centrifuged at 10,000 g for 20 minutes at 4°C. The supernatant was then concentrated, buffer exchanged with 10 kDa Amicon filters to 6 M urea, 50 mM sodium borate buffer, pH 8.2 and portion containing the 32 kDa LOX was collected. The presence of LOX was verified by means of Western Blot (refer to Fig. 1A).

**Fig 1.**
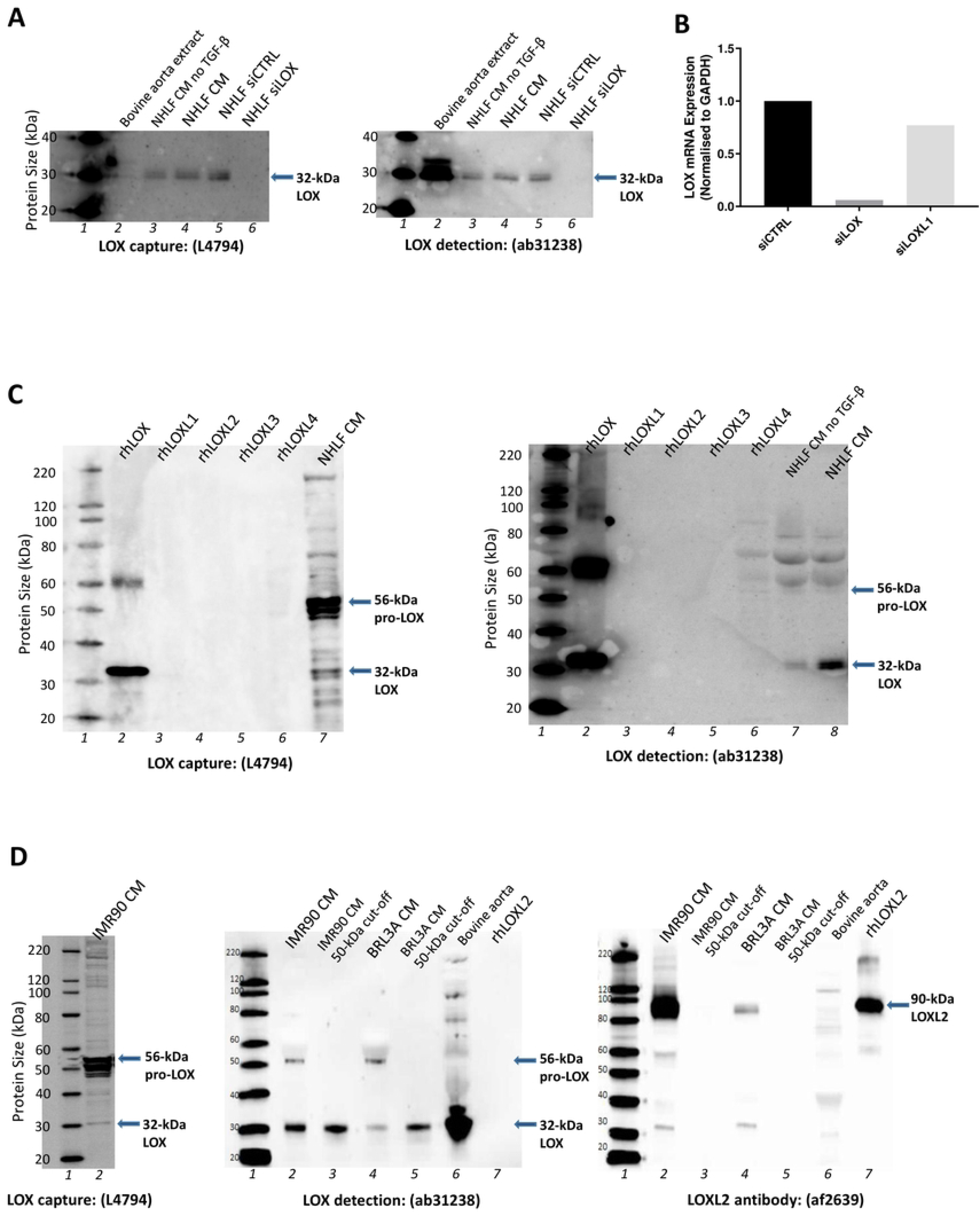
Validation of LOX antibodies for digital ELISA platform. **(A)** Evaluate specificity of the LOX antibodies using NHLF cells that underwent LOX siRNA knockdown. Lane 1: molecular weights ladder, with adjacent labels. Lane 2: LOX extracted from bovine aorta. Lane 3: protein derived from NHLF fibroblast cell condition medium (CM) without TGFβ supplement after mock transfection. Lane 4: protein derived from NHLF fibroblast CM after mock transfection. Lane 5: protein derived from NHLF CM after siRNA control (siCTRL) transfection. Lane 6: protein derived from NHLF CM after LOX siRNA (siLOX) transfection. *Left blot —* primary antibody (Sigma, L4794) and anti-rabbit secondary antibody (Abcam, ab6721); *right blot —* primary antibody (Abcam, ab31238) and anti-rabbit secondary antibody (Abcam, ab6721). **(B)** Confirmation of LOX siRNA knockdown in NHLF cells with RT-qPCR analysis of LOX mRNA expression. **(C)** Evaluate the selectivity of LOX antibodies. Lane 1: molecular weights ladder, with adjacent labels. Lane 2: recombinant human LOX. Lane 3: recombinant human LOXL1. Lane 4: recombinant human LOXL2. Lane 5: recombinant human LOXL3. Lane 6: recombinant human LOXL4. Lane 7: protein derived from NHLF fibroblast cell condition medium (CM) without TGFβ supplement after mock transfection. Lane 8: protein derived from NHLF fibroblast CM after mock transfection. *Left blot —* primary antibody (Sigma, L4794) and anti-rabbit secondary antibody (Abcam, ab6721); *right blot —* primary antibody (Abcam, ab31238) and anti-rabbit secondary antibody (Abcam, ab6721). **(D)** Evaluate the characteristics of LOX antibodies for pro-LOX and active LOX. *Left blot —* Lane 1: molecular weights ladder with adjacent labels. Lane 2: protein derived from full IMR90 CM. Primary antibody (Sigma, L4794) and anti-rabbit secondary antibody (Abcam, ab6721). *Middle and right blots —* Lane 1: molecular weights ladder with adjacent labels. Lane 2: protein derived from full IMR90 human fibroblast CM. Lane 3: protein from IMR90 CM with size above 50-kDa being excluded. Lane 4: protein derived from full BRL3A rat fibroblast CM. Lane 5: protein from BRL3A rat with size above 50kDa being excluded. Lane 6: LOX extracted from bovine aorta. Lane 7: recombinant human LOXL2. Middle: primary antibody (Abcam, ab31238) and goat anti-rabbit secondary antibody (Abcam, ab6721); right: primary antibody (RnD System, af2639) and anti-goat secondary antibody (Abcam, ab6741).

#### Recombinant human LOXL1 production

NIH-3T3 cells (ATCC) were transfected with pcDNA3.3.hLOXL1-Neo (Thermo Fisher Scientific, K830001) using Lipofectamine-LTX (Thermo Fisher Scientific, 15338500) and placed under geneticin (500 µg/mL, Thermo Fisher Scientific, 10131035) selection for 48 hours to produce stable expression of human recombinant LOXL1. The cells stably expressing human recombinant LOXL1 were maintained in culture (37°C, 5% CO_2_) at log-phase growth in advanced DMEM (Thermo Fisher Scientific, 12491015) with 10% heat-inactivated fetal bovine serum (FBS, Thermo Fisher Scientific, A3161001) and geneticin. At 90% confluency, cells were incubated in starving medium containing advanced DMEM with 2.5% heat-inactivated FBS for 3 days. The CM was subsequently collected, filtered (0.45 µM), and the His-tagged LOXL1 was column purified by means of Ni^2+^-Sepharose 6 Fast Flow and His buffer kit (Cytiva, 17-5318-06) as per manufacturer’s instructions. The elution run-through was concentrated, buffer exchanged with 10 kDa Amicon filters to buffer containing 1.2 M urea, 50 mM sodium borate, pH 8.2, and the portion containing the ∼ 63 kDa LOXL1 was collected.

#### Recombinant human LOXL4 production

HEK293 cells stably expressing human recombinant LOXL4 were generated and provided by Dr. Fernando Rodríguez-Pascual (C.S.I.C., Universidad Autónoma de Madrid, Madrid, Spain, as detailed (14)). Cells were cultured (37°C, 5% CO_2_) in growth medium containing DMEM (Thermo Fisher Scientific, 11965092) with 10% Tet System Approved heat-inactivated FBS (Takara Bio; 631101), 200 µg/mL hygromycin (Thermo Fisher Scientific, 10687010) and NEAA (Thermo Fisher Scientific, 11140050). At 90% confluency, cells were washed twice with warm sterile PBS and incubated (37°C, 5% CO_2_) in starving medium containing DMEM phenol red free with 10 µM CuSO_4_. Doxycycline (1 µg/mL, Sigma-Aldrich, D3072) was added to induce recombinant protein expression. After 2 days, the CM was collected, filtered (0.45 µM), concentrated and buffer exchanged with 50 kDa Amicon filters to buffer containing 1.2 M urea, 50 mM sodium borate, pH 8.2, and the portion containing the ∼ 82 kDa LOXL4 was collected.

#### Other recombinant human lysyl oxidases

Recombinant human (rh) LOX was purchased from either LSBio (LS-G3047 or LS-G22712) or Origene (TP313323). None of these rhLOX were enzymatically active in our assays. rhLOXL2 (2639-AO) and rhLOXL3 (6069-AO) were enzymatically active, purchased from R&D Systems.

#### Human skin LOX extraction

To extract LOX from the 3 mm human skin biopsy, epidermal and dermal layers of the skin were dissected, pre-cooled with liquid nitrogen and pulverized. Samples were washed thrice by homogenization with ice-cold wash buffer (0.15 M NaCl, 50 mM sodium borate, pH 8.0 with 0.25 mM PMSF and 1 µL/mL bovine aprotinin as protease inhibitors) at 100 µL/mg, centrifuged at 10,000 *g* for 10 minutes at 4°C and supernatant discarded. After the final washing step, the pellet was resuspended in extraction buffer (6 M urea, 50 mM sodium borate, pH 8.2 with 0.25 mM PMSF and 1 µL/mL bovine aprotinin) with ratio of buffer volume to tissue weight at 4:1. After 3-hour incubation at 4°C, the mixture was diluted 1:2 with assay buffer containing 50 mM sodium borate (pH 8.2), centrifuged at 20000 g for 20 minutes at 4°C. The collected supernatant was spiked with pargyline hydrochloride (Sigma-Aldrich, P8013) and mofegiline hydrochloride (Sigma-Aldrich, ATE245843861) at final concentrations of 0.5 mM and 1 µM, respectively, to inhibit other amine oxidases. LOX concentration in the final supernatant was determined by DEA. LOX enzymatic activity in the skin extract was determined using a fluorescent based biochemical assay where LOX activity was monitored by H_2_O_2_ generated by LOX through the horseradish peroxidase (HRP)-catalyzed conversion of Amplex red to fluorescent resorufin (12).

### Extracellular vesicle isolation

Two batches of IMR-90 fresh CM were generated as described in the above section – “Native LOX production and extraction from fibroblast cell lines”. Isolation of the extracellular vesicle (EV) has been described previously (15). Briefly, the fresh CM was centrifuged at 10,000 g for 30 minutes to precipitate large EVs. For isolation of nano-sized EVs, the supernatant remaining after the 10,000 *g* spin was subjected to additional centrifugation performed on a 30% sucrose cushion (or without sucrose) at 100,000 *g* for 1 hour at 4°C. The resulting EV pellets were washed with PBS (1 mL) and stored at −80°C until further analysis. The remaining supernatant was further concentrated (∼ 40×) and buffer exchanged to 6 M urea, 50 mM sodium borate buffer, pH 8.2 with 10 kDa Amicon filters. The EVs and supernatants were analysed in Western blot for specific EV markers, and LOX expression was determined by either Western blot or DEA.

### Lysyl oxidase knockdown

#### siRNA transfection

The NHLF cell line (Lonza, CC-2512B) was cultured in its respective growth medium as per manufacturer’s instructions supplemented with or without 10 ng/mL rhTGF-β, in an atmosphere consisting of 5% CO_2_ in air at 37°C in a humidified incubator. ON-TARGETplus Smart pool small interfering RNA (siRNA) duplexes were used: scrambled non targeting control (siCTRL) (Dharmacon, D-001810-10-05) or for LOX (Dharmacon, L-009810-00-0005)-or LOXL1 (Dharmacon, L-004645-01-0005) knockdown. The siRNAs were diluted in buffer according to the manufacturer’s instructions. NHLF cells were seeded in duplicate in six-well plates with reverse transfection conditions as described by the Manufacture. As previously mentioned the LOX or LOXL1 siRNA or siCTRL were complexed with the transfection reagent Lipofectamine RNAiMax (Thermo Fisher Scientific, 13778075) and added to the NHLF cells. siRNAs for both LOX and LOXL1 were used at a concentration of 50 nM and incubated overnight. The media was replaced and the cells were allowed to recover for 24 hrs. The cells were washed with sterile PBS and incubated in starving medium with or without 10 ng/mL rhTGF-β for 48 hrs. The cell lysates were collected and evaluated for LOX knockdown by RT-qPCR, while the concentrated CM was evaluated for LOX knockdown by Western blotting.

#### cDNA synthesis and quantitative real time PCR

The RNA extraction (miRNeasy Kit) and on-column DNasI treatment were performed, as per the manufacturer’s instructions (Qiagen, 217004). The RNA concentration was evaluated by measuring the absorbance at 260 and 280 nm using NanoDrop^TM^ Spectrophotometer (Thermo Fisher Scientific). A conventional reverse transcription reaction was performed to yield single-stranded cDNA using SuperScript VILO cDNA Synthesis Kit (Thermo Fisher Scientific, 11754050). The PCR cycling parameters were 15 seconds × 40 cycles at 95°C. Glyceraldehyde-3-phosphate dehydrogenase (GAPDH) was used as the house keeping gene in comparison of gene expression data. Gene expression was measured by the 2^−ΔCt^ method using ABI7500 (Applied Biosystems) with the ABI TaqMan primer sets as specified: *LOX* (Hs00942483_m1); *LOXL1* (Hs00935937_m1); *Gapdh* (Hs02786624_g1).

### Lysyl oxidase activity assay

#### Fluorescent-based biochemical assay

For the detection of lysyl oxidase activity in the IMR90 CM (full CM or 50-kDa cut-off CM) and the human skin biopsy extract, the mix was spiked with pargyline hydrochloride and mofegiline hydrochloride at final concentrations of 0.5 mM and 1 µM, respectively, to inhibit other amine oxidases. The samples (25 µL) were then incubated with either 100 nM PXS-5120 (a selective LOXL2 inhibitor with similar pharmacological profiles to PXS-5153A (16), applicable to IMR90 CM sample only to determine specific enzymatic activity of LOX versus LOXL2), 600 µM β-aminopropionitrile (BAPN, a pan-lysyl oxidase inhibitor) or DMSO (control) for 30 minutes at 37°C and assayed by adding 25 µL of reaction mixture containing 50 mM sodium borate, 120 µM Amplex Red, 1.5 U/mL horseradish peroxidase, and 20 mM putrescine dihydrochloride as substrate, pH 8.2. Resorufin fluorescent signal intensity was measured every 2.5 min for 30 minutes at 37°C (with excitation and emission at 544 nm and 590 nm, respectively) and the enzymatic activity was indicated by the rate of the signal (kinetic value without BAPN).

#### Activity-based probe assay

LOX activity in the human serum was measured using an activity-based probe for the detection of enzyme activity in conjunction with the single molecule array (SIMOA) bead technology as previously described (13).

### Western blot

Recombinant human (rh) LOX (LSBio, LS-G3047), rhLOXL1-4, bovine LOX extract, IMR90 EVs and concentrated CM from various cell lines were denatured at 70°C for 10 minutes and run on NuPAGE® Novex® 4-12% Bis-Tris gel system (Invitrogen, NP0321BOX). Gels were electroblotted onto the nitrocellulose membrane (Amersham Pharmacia Biotech), and target proteins were captured by the corresponding primary-secondary antibody pairs. Primary antibodies used were: LOX capture (Sigma-Aldrich, L4794, 1:500), LOX detection (Abcam, ab31238, 1:500), LOXL2 (R&D Systems, AF2639, 1:500), rabbit anti-human LOXL4 (provided by Fernando Rodriguez-Pascual, 1:500), Alix (Cell Signaling, E6P9B, 1:1000) and CD81 (Cell Signaling, D3N2D, 1:1000). Secondary antibodies used were goat anti-rabbit (Abcam, ab6721, 1:5000 or 1:10000) or rabbit anti-goat (Abcam, ab6741, 1:5000 or 1:10000). Super-signal West Pico Chemiluminescent Substrate (Thermo Fisher Scientific, PI-34087) was used for signal development and ChemiDoc™ Touch Imaging System (NRNQ1432) was used for imaging.

### SIMOA – a component of DEA

The DEA platform consists of the Simoa^TM^ (single molecule array) bead technology to isolate the protein in conjunction with an antibody for the detection of total protein. This platform has enabled accurate measurement of total LOX protein from small volumes of serum, plasma and tissue (S1 Fig). In the immunoassay, paramagnetic capture beads are coated with a LOX capture antibody (Sigma-Aldrich, L4794). 100 µL of pre-diluted (4-fold) sample in buffer (0.5% Casein, 0.25% Tween-20 in PBS) is combined in a cuvette with the capture beads (25 µL for protein content at 1.6 x 10^7^ beads/mL). The biotinylated detector agent, a second anti-LOX antibody (Abcam, ab219369), is then added during the same incubation with intermittent shaking at room temperature for 35 minutes. Target molecules present in the sample are captured by the antibody coated beads and bound with the detector agent simultaneously. After incubation, beads are pelleted by magnetic separation, excess sample/buffer and reagents aspirated off and beads resuspended in wash buffer to remove unbound proteins and excess reagents. Following protein capture and incubation with the detector, a conjugate of streptavidin-β-galactosidase (SβG; 100 µL at 300-350 pM) is mixed with the beads and incubated for 5 minutes with intermittent shaking at room temperature. SβG binds to the biotinylated detector agent, resulting in enzyme-labelling of captured LOX. Following a final wash, the beads are resuspended in a resorufin-β-D-galactopyranoside (RGP) substrate solution and transferred to a SimoaTM Disc (containing a microarray to separate beads into individual microwells). After settling into the microarray via gravity, beads are then sealed in microwells with oil. If LOX is captured and labelled on the bead, β-galactosidase hydrolyses the RGP substrate in the microwell into a fluorescent product that provides the signal for measurement. A single-labelled target molecule results in sufficient fluorescent signal that can be detected and counted by the SimoaTM optical system. The percentage of bead-containing wells in the array that have a positive signal is proportional to the amount of target present in the sample. To determine LOX protein concentration, data can be interpolated from a calibration curve using rhLOX (LSBio™, LS-G3047). All samples are measured in triplicate and the mean is presented per sample.

Note the LOX detection antibody used in the DEA (Abcam, 219369) is derived from the same clone as the antibody tested in the western blot optimization (Abcam, 31238) but without the carrier protein.

### Commercial ELISA kits

A human competitive LOX ELISA kit was tested (MyBioSource, MBS745192, lot BG190318HUC). Assays were performed twice on the same plasma samples as per manufacturer’s instructions.

A human sandwich LOX ELISA kit was tested (Cloud-Clone, SEC580Hu, lot L180817140). Assay was performed twice on the same plasma samples (as above) as per manufacturer’s instructions.

### Analysis

Data was presented as mean ± SD. Statistics were analysed by Graphpad Prism v8.4. For the DEA calibration curve, a linear regression was performed between the signal units and rhLOX. Comparisons between the LOX values of bladder cancer patients and the healthy subjects were performed using an unpaired t-test. Pearson correlation was performed between LOX activity and LOX concentration in serum and skin extract. Pearson correlation was also performed between averaged plasma concentrations from the two competitive ELISA tests and the concentration derived from the DEA.

## Results

Due to the variability of existing ELISA kits, and the need for a consistent assay for the detection of LOX in human clinical samples, we assessed commercially available LOX antibodies for their suitability in a sandwich ELISA, which could subsequently be transferred to the SIMOA platform.

### Antibody specificity and selectivity

The inaugural step in the development of a sensitive digital ELISA for LOX was assessing the specificity and selectivity of the antibodies binding to active human LOX. NHLF cells secrete large amounts of LOX enzyme; we utilized siRNA to assist in determining the specificity of the antibodies. The concentrated CM from NHLF cells was tested for LOX activity and concentration. Both antibodies detected the 32 kDa LOX band in the CM of the control NHLF cells (Fig 1A, Lane 4) and in cells that underwent control knockdown (siCTRL, Fig 1A, Lane 5). In contrast, cells that underwent specific LOX knockdown (siLOX) showed complete ablation of LOX expression (Fig 1A, Lane 6). The knockdown was also confirmed by the quantitative PCR analysis of the cell lysate, showing nearly-complete reduction of LOX mRNA expression in the siLOX cells. For comparison, LOX expression was not affected in those cells that underwent LOXL1 knockdown (Fig 1B).

The selectivity of the two LOX antibodies was evaluated against the recombinant human lysyl oxidase family members. Both antibodies detected rhLOX at 32 kDa; the band at ∼ 60 kDa (Fig 1C Lane 2 of the left and right blots) appeared to be the dimer of rhLOX. None of the two LOX antibodies bound to rhLOXL1-4, demonstrating their selectivity for rhLOX (Fig 1C, Lanes 3-6 of the left and right blots). As rhLOXL1 and rhLOXL4 were produced by non-commercial sources, they were further verified by Western blot as shown in S2 Fig.

Specificity of the LOX antibodies for native human LOX was confirmed: both antibodies detected the 32 kDa active LOX and the 56 kDa pro-LOX in the CM of the NHLF (Fig 1C, Lane 7 of the left blot and Lane 8 of the right blot) and IMR90 (Fig 1D, Lane 2 of the left and middle blots) cell lines. IMR90 CM was also used to confirm the selectivity of the LOX antibodies over native LOXL2. The 90 kDa native LOXL2 (in the IMR90 full CM) and the rhLOXL2, detected by the anti-LOXL2 antibody (Fig 1D, Lane 2 and 7 of the right blot), were not shown in the LOX blot (Fig 1D, Lane 2 and 7 of the middle blot), demonstrating the LOX detection antibody was selective for native LOX over LOXL2. Similarly, antibody selectivity was shown for the native rat LOX over LOXL2 using the BRL3A CM (Fig 1D, Lane 4 and 5 of the middle blot).

### DEA setup and sensitivity

The 32 kDa LOX from IMR90 CM (50 kDa cut-off) was used to optimize the conditions for DEA and to test the assay sensitivity, given LOX is the only member of the family with a molecular weight < 50 kDa. Native LOX from IMR90 CM has the same molecular size as rhLOX (Fig 2A), but, in contrast to inactive rhLOX, was enzymatically active. This was demonstrated by a complete inhibition of activity in the presence of a pan-LOX inhibitor (BAPN) but not by the selective LOXL2 inhibitor (PXS-5120). In contrast, the full IMR90 CM (with a mixture of lysyl oxidase isoforms) was sensitive to a selective LOXL2 inhibitor with ∼ 50% activity being diminished (Fig 2B) in line with the data from Western blots.

**Fig 2.**
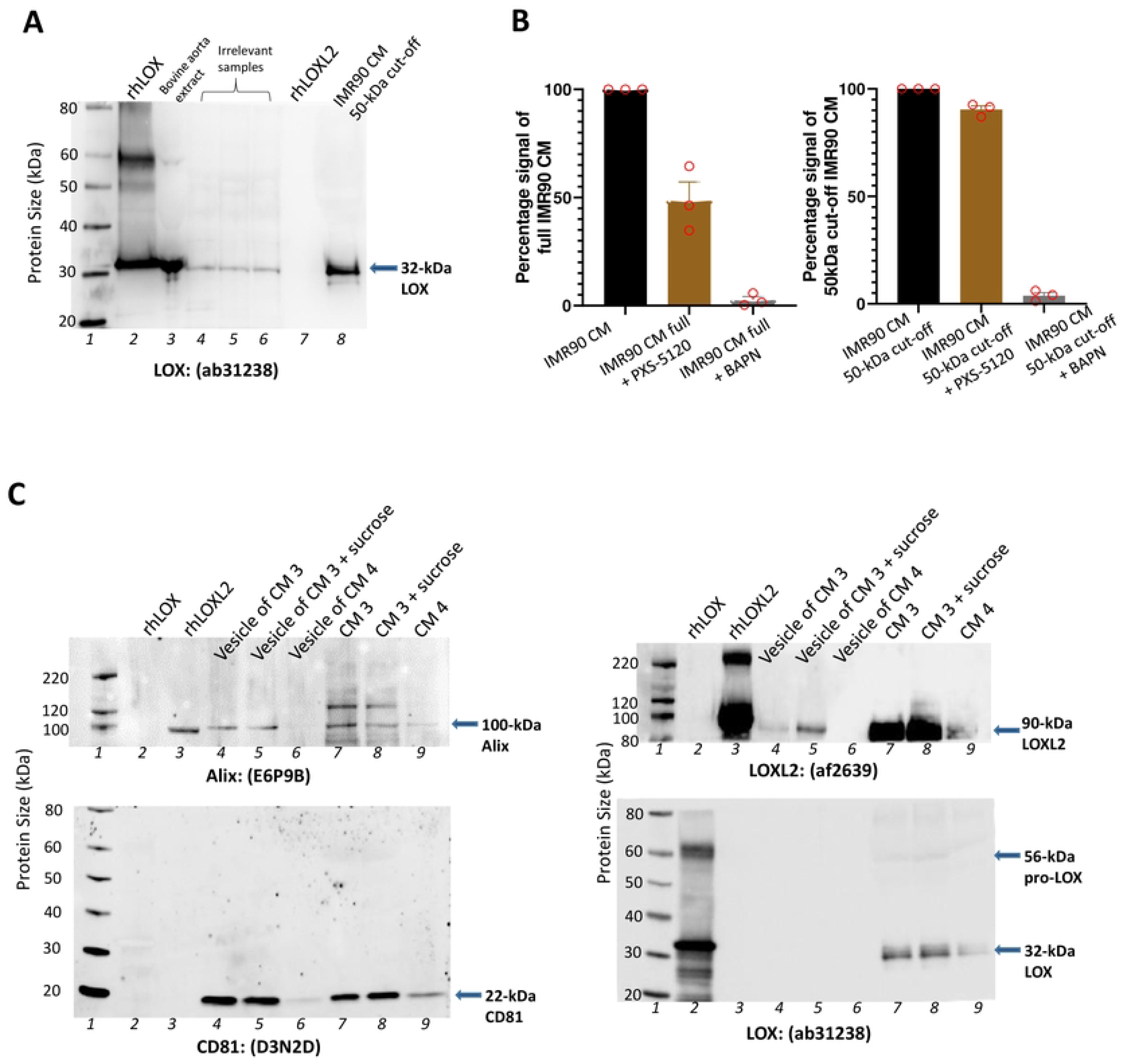
Validation of digital ELISA platform with the measurement of native LOX from human fibroblast cell line IMR90. **(A)** Western blot validating the native LOX expression in IMR90 cell conditioned medium (CM). Lane 1: molecular weights ladder, with adjacent labels. Lane 2: recombinant human LOX. Lane 3: LOX extracted from bovine aorta. Lanes 4-6: animal tissue samples (irrelevant to current study). Lane 7: recombinant human LOXL2. Lane 8: protein derived from IMR90 CM with size above 50 kDa being excluded. Primary antibody (Abcam, ab31238) and goat anti-rabbit secondary antibody (Abcam, ab6721). **(B)** Lysyl oxidases activity in full IMR-90 CM or IMR90 CM with size above 50 kDa being excluded, analysed by measuring fluorescent signal generated with amplex red (30 µM), HRP (0.75 U/mL) and the lysyl oxidase substrate putrescine (10 mM). A selective LOXL2 inhibitor PXS-5120 (100 nM) or a pan-lysyl oxidase inhibitor β-aminopropionitrile (BAPN, 100 µM) were used in the assay to determine the specific enzymatic activity. **(C)** Western blot showing LOX expression in extracellular vesicles isolated from IMR90 CM. Lane 1: molecular weights ladder, with adjacent labels. Lane 2: recombinant human LOX. Lane 3: recombinant human LOXL2. Lane 4 and 6: extracellular vesicles isolated from full IMR90 CM. Lane 5: extracellular vesicles isolated from full IMR90 CM supplemented with sucrose. Lane 7 and 9: full IMR90 CM. Lane 8: Full IMR90 CM supplemented with sucrose. Left top blot—primary anti-human Alix antibody (Cell Signaling Technology, E6P9B) and goat anti-rabbit secondary antibody (Abcam, ab6721). Left bottom blot— primary anti-human CD81 (Cell Signaling Technology, D3N2D) and goat anti-rabbit secondary antibody (Abcam, ab6721). Right top blot— primary antibody (RnD System, af2639) and rabbit anti-goat secondary antibody (Abcam, ab6741). Right bottom blot— primary antibody (Abcam, ab31238) and goat anti-rabbit secondary antibody (Abcam, ab6721).

In order to optimize the signal for the LOX ELISA, both LOX antibodies were tested as either the capture or the detection antibody, using IMR90 50 kDa cut off LOX spiked into buffer or plasma/serum. The signal to noise was larger with the L4794 for capture than ab31238. A carrier protein free LOX antibody of ab31238 (Abcam, ab219369) was used in DEA as the detection antibody to further lower background signal (S3 Fig).

Known concentrations of rhLOX were used to establish a calibration curve for DEA. A linear curve was generated by plotting serial dilutions (from 100 down to 0.02 ng/mL) of the rhLOX protein against the assay signal generated from each concentration (S4 Fig). The plot demonstrated an assay sensitivity of ∼0.1 ng/mL. Native protein was used to demonstrate assay robustness when diluting samples. Using the 50 kDa cut-off IMR90 CM in different dilutions (IMR90 CM 1 and 2, Table 1), the difference between the corrected LOX value of 10x different samples was less than 10%.

**Table 1.** LOX concentration in IMR90 cell conditioned medium and extracellular vesicle fraction measured with DEA.

| Sample | Digital ELISA (DEA) |  |  |
| --- | --- | --- | --- |
| | Dilution applied | Conc. measured (ng/mL) ± variation in replicates (%) | Corrected conc. (ng/mL) |
| IMR90 CM 1<br>50 kDa cut-off | 1:500 | 0.54 (± 2%) | 270 |
| | 1:50 | 6.0 (± 2%) | 300 |
| IMR90 CM 2<br>50 kDa cut-off | 1:100 | 0.10 (± 5%) | 10 |
| | 1:50 | 0.21 (± 5%) | 11 |
| IMR90 CM 3 | 1:50 | 0.80 (± 3%) | 40 |
| IMR90 CM 3 +<br>sucrose | 1:50 | 0.59 (± 0%) | 30 |
| IMR90 CM 4 | 1:50 | 0.42 (± 1%) | 21 |
| Vesicle of IMR90<br>CM 3 | 1:50 | 0.0082 (± 7%) | 0.41 |
| Vesicle of IMR90<br>CM 3 + sucrose | 1:50 | 0.018 (± 2%) | 0.91 |
| Vesicle of IMR90<br>CM 4 | 1:50 | 0.0055 (± 15%) | 0.025 |

To demonstrate the superiority of DEA over conventional Western blots, LOX concentration in the extracellular vesicles (EVs) extracted from IMR90 CM was measured. EV pellet showed vesicle-specific markers Alix and CD81, confirming successful EV extraction (Lanes 4-6 of the left top and bottom blots, Fig 2C). In the CM, Alix and CD81 were positivity displayed at a similar level to the vesicles (Lanes 7-9 of the left top and bottom blots, Fig 2C). While both LOX and LOXL2 proteins can be detected using western blot in the CM (Lanes 7-9 of the right top and bottom blots, Fig 2C), the LOX capture antibody failed to demonstrate LOX expression in the EVs when using Western blots (Lanes 4-6 of the right bottom blot, Fig 2C). In contrast, LOXL2 protein was measurable in the same assay format using the same EVs (Lanes 4-5 of the right top blot, Fig 2C). As expected, LOX is present in the EV extraction using DEA which enables to the detection of LOX at lower levels. The LOX concentration detected in each of the EV fraction was 30-100 folds lower compared to the CM it was extracted from (Table 1).

### Application of DEA

#### Human serum

Using the optimized LOX DEA, we investigated the concentration of the LOX enzyme in serum. Interestingly, in serum, bladder cancer patients had significantly higher LOX concentrations compared to matched healthy subjects (Fig 3A). Furthermore, the LOX concentration correlated with the LOX activity (measured by the activity-based bioprobe assay (13)) (Fig 3B) providing an independent validation of DEA.

**Figure 3.**
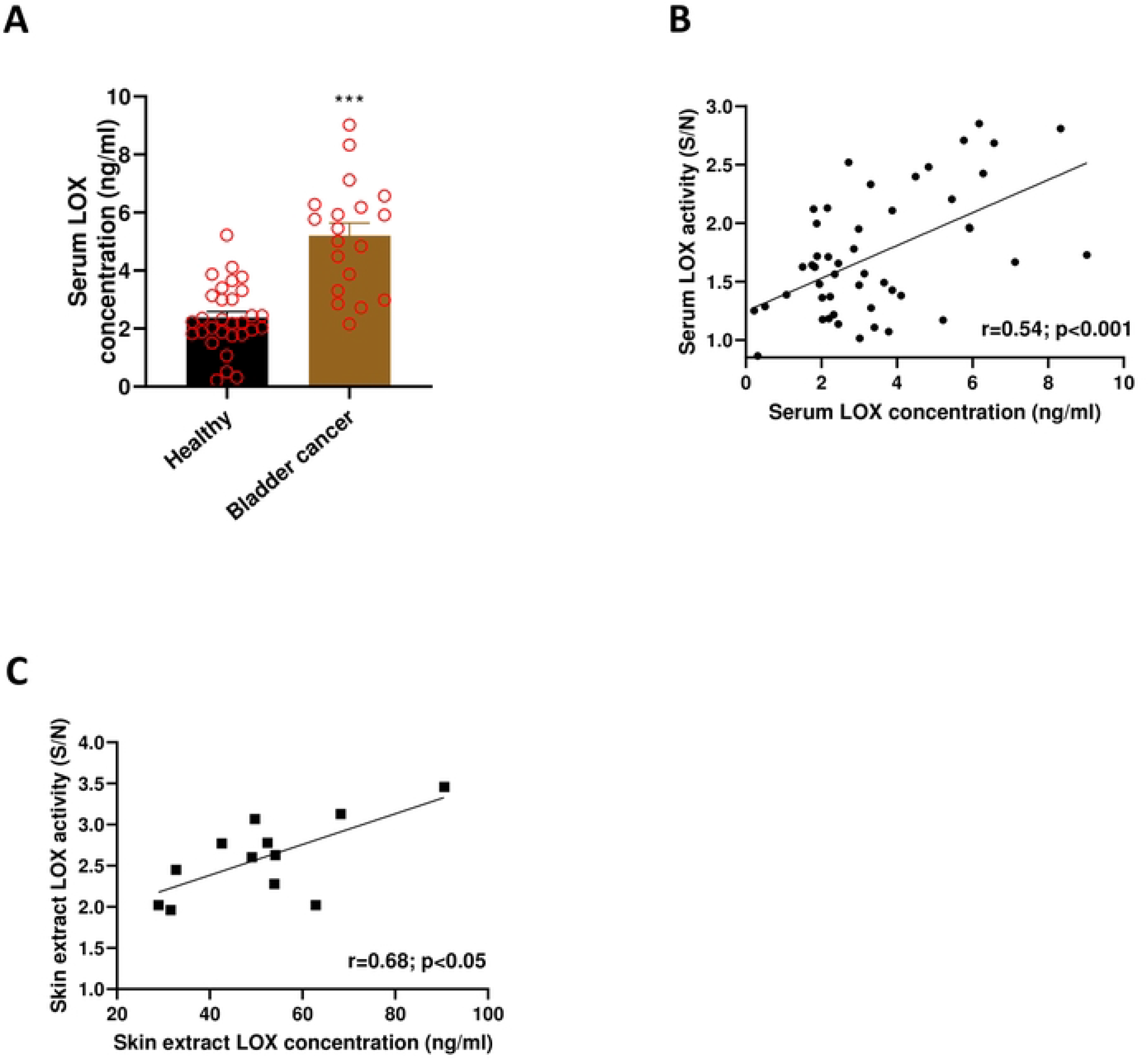
LOX concentration in human serum and skin tissue extract measured by DEA. **(A)** Serum LOX concentration in healthy subjects and bladder cancer patients measured with digital ELISA assay (DEA). Data presented as mean ± SD. *** P<0.001 vs healthy group. **(B)** Correlation between LOX activity measured by activity-based probe assay and LOX concentration measured by DEA in human serum samples acquired from healthy and bladder cancer patients. **(C)** Correlation between LOX activity measured by the fluorescent based biochemical assay [amplex red (30 µM), HRP (0.75 U/mL) and the lysyl oxidase substrate putrescine (10 mM)] and LOX concentration measured by DEA in skin biopsy extracts acquired from healthy human subjects.

#### Human skin extract

We next sought to determine whether the DEA was able to quantify LOX in small tissue extracts (skin biopsies). Three-millimeter cross sectional skin biopsies were obtained from healthy human subjects and LOX was extracted according to methods outlined. LOX concentration from each skin extract was measured by DEA and LOX activity analyzed using the fluorescent-based biochemical assay. It was determined that the LOX concentration significantly correlated with activity (Fig 3C). These data confirmed that our novel DEA can accurately detect LOX concentrations isolated from both liquid biopsies and tissue extracts.

#### DEA versus commercial ELISAs

The sensitivity and accuracy of DEA was further compared with commercial ELISA kits for measuring human LOX concentration in human plasma. In the plasma samples of eight healthy subjects, the LOX concentration measured using the commercial competitive ELISA returned a much lower value (10-25 fold) compared to DEA (S5 Fig and S2 Table). It was interesting to note, that the values obtained from the two assays were correlated (S6 Fig), indicating that the competitive ELISA, despite underestimating the absolute values, was able to resolve the difference between samples. Another commercial sandwich ELISA kit for human LOX was also tested but failed to detect LOX-specific signals in IMR90 CM or human plasma (data not shown).

## Discussion

The current study illustrates that the DEA for LOX is a valid, sensitive and accurate assay for determining the absolute LOX concentrations in human samples from both blood and tissue.

Commercially available rhLOX has been shown to have little to no enzymatic activity, possibly due to the lack of glycosylation of the pro-region involved in the correct folding of the mature protein (17). In order to test the sensitivity and accuracy of the DEA system for the LOX protein, the 32 kDa LOX produced from IMR90 cells, which we have demonstrated functional activity, was used in the current study.

The capture (L4797, Sigma-Aldrich) and detection (ab31238, Abcam) LOX antibodies in the study were chosen due to their superior specificity and selectivity against the recombinant and native human LOX proteins in addition to the lack of interference. The capture antibody that detects both the 32 kDa LOX and the 56 kDa pro-LOX could capture all fragments of LOX in a given sample. As shown in the western blots, the detection antibody has a higher affinity for the active 32 kDa LOX that catalyzes the crosslinking process, over the 56 kDa pro-LOX that is known to be enzymatically inactive. Consequently, the measured value will be more reflective of the functional relevant active LOX in the physiological system.

The concentrations of LOX measured by the DEA have shown to be significantly correlated to the LOX enzymatic activity in the human serum and skin tissue extract. Any correlative deviation between the concentration assay and the functional assay is likely due to the contribution of the enzymatically inactive 56 kDa form in DEA which is not measured in the functional assay. Additionally, inactive smaller LOX fragments may be also captured in DEA.

The current DEA detected very low expression levels of LOX in the EVs, which was not detectable by the conventional Western blot. It is known that LOXL2 was expressed in the EVs of endothelial (18) and cancerous cell lines (19), in contrast no reports to date indicate the presence of LOX in EVs, potentially due to its low concentration.

In conclusion, these data demonstrate that a specific, selective and accurate measurement of LOX in small, easily accessible, human samples is possible. Additionally, they also show the need for thorough evaluation of available ELISA kits to ensure accuracy. The highly sensitive and accurate DEA for LOX is suitable to be used in the clinic to investigate the role of LOX as a diagnostic in fibrosis and cancer.

## Acknowledgements

The authors would like to thank Heidi Schilter for her inputs in the initial phase of the experimental design. The authors would also like to thank the Quanterix Team for running the DEA assays.

## Author contributions

Y.Y., L.P. and W.J. involved in conception and design. Y.Y., L.P., A.Z., R.H. and J.S. involved in acquisition and analysis. All authors involved in drafting and revising the manuscript. All authors approved the final manuscript and agreed to be accountable for all aspects of the work.

